# Elucidating the mechanism of microbial thermogenesis

**DOI:** 10.1101/2023.06.02.543367

**Authors:** Puneet Singh Dhatt, Tae Seok Moon

## Abstract

Organisms necessarily release heat energy in their pursuit of survival. This process is known as cellular thermogenesis and is implicated in many processes from cancer metabolism to spontaneous farm fires^1,2^. However, the molecular basis for this fundamental phenomenon is yet to be elucidated. Here, we show that the major players involved in cellular thermogenesis are the protein kinases ArcB, GlnL, and YccC in *Escherichia coli*. We also reveal the substrate-level control of ATP-driven autophosphorylation that governs cellular thermogenesis. Through live-cell microcalorimetric experimentation, we find that only three of the 231 regulatory proteins, when knocked out in a model *Escherichia coli* strain, dysregulate cellular thermogenesis. This dysregulation can be seen in an average 25% or greater increase in heat output by these cells. We also discover that both heat output and intracellular ATP levels are maximal during the late log phase of growth. Our results demonstrate a correlation between ATP concentrations in the cell and a cell’s ability to generate excess heat. We expect this work to be the foundation for engineering a new generation of thermogenically-tuned organisms for a variety of applications.

---

The concepts of life and heat have been considered interdependent since the time of the ancients^3^. Aristotle was the first person in recorded history who formed a theory on the origin of life. In his treatise *Generation of Animals* in the 4^th^ century BCE, he wrote on the importance of a “vital heat” that is responsible for animating organic material into life^4^. Yet, there still remains the underexplained fact that all organisms release heat^5^.

Studies on cellular thermogenesis became popular at the end of the 18^th^ century after the first calorimeters were constructed to measure live cell heat generation^6–10^. Adair Crawford’s seminal work in 1779 studied the heat output of guinea pigs^11^. Just one year later, Lavoisier and Laplace demonstrated the connection between metabolism and thermogenesis using a novel ice-based calorimeter^12,13^. To this day, the leading theory in molecular biology is that heat is generated at the molecular level through metabolism. In fact, it is said that 50-60% of the energy that is stored in metabolic substrates is released as heat^14^.

At the beginning of the 20^th^ century, work in cellular thermogenesis was largely focused on studying microorganisms’ ability to cause “spontaneous heating” of farm products. A process which, to this day, causes farm fires across the world^2,15–22^. This work went on to influence cancer research with the establishment of the Warburg Hypothesis in 1924^1,23^. Recent work in microbial thermogenesis is mainly focused on the use of isothermal microcalorimetry to monitor microbial activities such as the metabolic dynamics of soil bacteria^24,25^, marine sedimental heat production^26^, and antibiotic resistance behaviors to determine both antibiotic mechanism and optimal dosage^27^. In eukaryotic systems, microcalorimetry has been applied to study cell thermogenesis to understand antiviral responses of pharmaceuticals^28,29^, study mitochondrial and endoplasmic reticular functions^30^, explain pathogen origins of obesity and anemia^30–33^, and explore cancer phenotypes for advanced diagnostics^34,35^.

Given that cellular thermogenesis is a fundamental property of organisms, we need to understand its impacts on complex symbiotic systems in nature. One such system is the human microbiome. It has recently been calculated that up to 70% of the energy released by a human each day can be attributed to the human microbiome^36^. Furthermore, microbial heat contribution has been observed in healthy animal models wherein antibiotic treatment lowers average body temperature by an average of 1°C^37–40^. This implies a key role of the human microbiome in human energy homeostasis. Gut flora have already been proven to be involved in numerous host regulation mechanisms, including endocrine function,^41^ genetic expression^42,43^, aging^44^, and immune processes^45^.

To our knowledge, the field of cellular thermogenesis has thus far been strictly observational. To elucidate the important metabolic agents and gain a fundamental mechanistic understanding of microbial thermogenesis, we hypothesized that there are a set of regulatory proteins that determine a cell’s thermogenesis. To test this hypothesis, we screened a single-gene knockout library of a model organism, *Escherichia coli* BW25113, for thermogenesis. The mechanistic understanding gained from this work can then be used to engineer new strains with altered thermogenesis and transition the field from observation to creation in the future.

In this work, we identify both a set of proteins that regulate microbial thermogenesis and the key metabolite, adenosine triphosphate (ATP), through which this regulation occurs in *E. coli*. A mathematical model was also developed to predict thermogenesis using ATP as the sole metabolite upon which thermogenesis is dependent. This work will allow us to engineer microbial thermogenesis to develop increased thermogenic strains for many applications in the future.

## Results and Discussion

### Experiment design using calorimetry

Calorimetric data is recorded using a thermocouple to correlate temperature changes to an electric signal (voltage) that is then calculated as heat energy by computer software. In general, there is a significant trade-off between the number of samples and sensitivity for calorimetry instruments, especially above eight sample wells^24^. Considering this trade-off, we chose a TAM Air Calorimeter, which has a sensitivity of approximately 1*μ*W. The initial seeding density of cells was optimized so that our final maximal signal for heat flow was reached at 5h. The baselining was done to settle for 12h before measurements were taken, such that the throughput of these experiments is heavily limited at just six experimental samples, along with one negative control and one positive control, tested per experiment. The experimental workflow is shown in Figure 1. Given this limitation in throughput, we tested rationally selected mutant groups of a single-gene knockout library^46^. We hypothesized that there is a global regulatory pathway for heat generation. In pursuit of this hypothesis, we tested all of the 199 transcription factors. We also included 29 histidine kinases (HKs) and 32 response regulators (RRs), which are often involved in global regulation through two-component signal transduction pathways^47^. The regulated pathways include carbon metabolism, nitrogen assimilation, and chemotaxis among others^48^. Furthermore, we tested 19 proteins related to central metabolism and 120 other proteins. There are 29 proteins that belong to both the transcription factor and RR groups, meaning 370 unique single-gene knockouts tested in this study.

**Figure 1:**
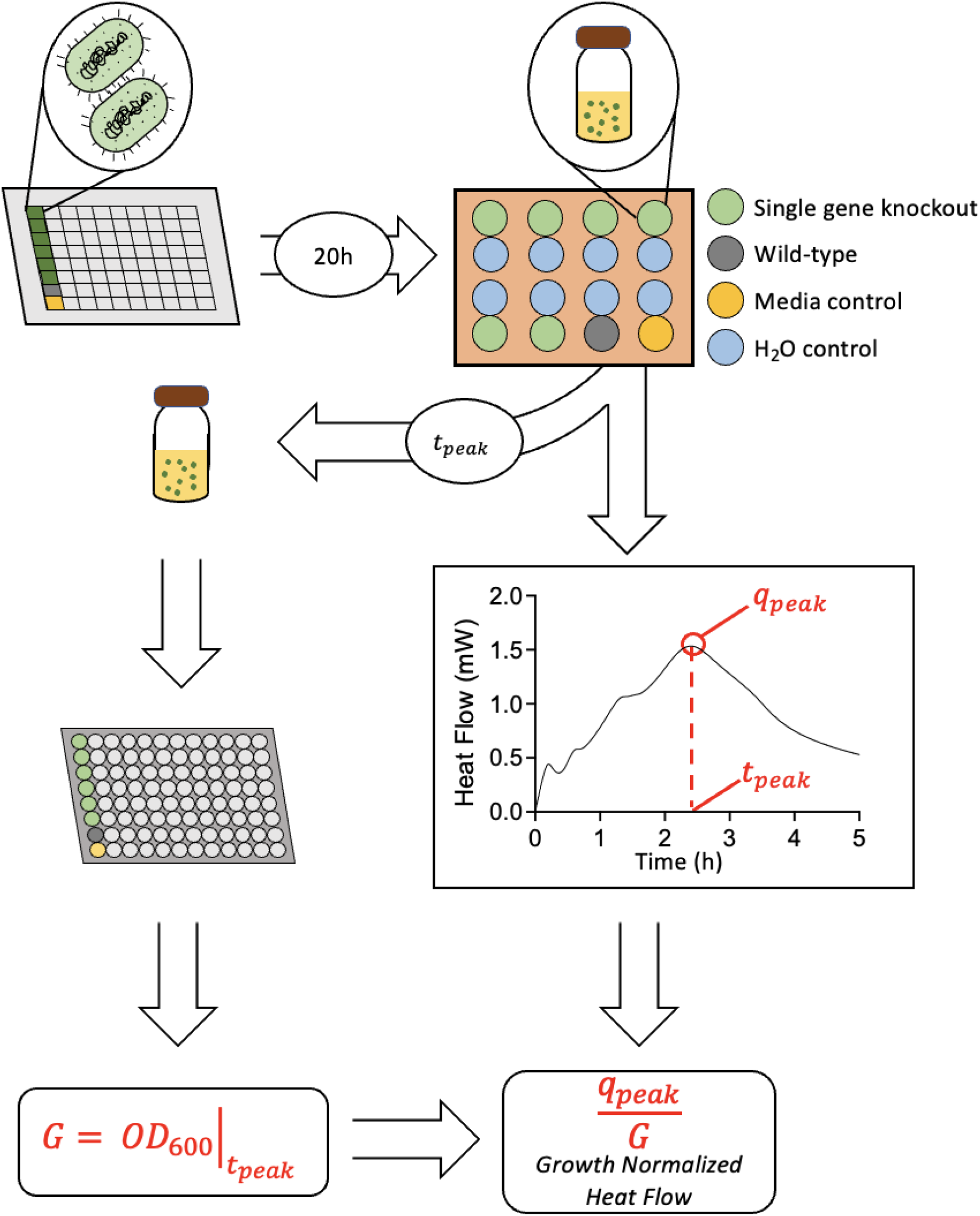
Methodology of live-cell microcalorimetry experiments. The method by which cells are grown, seeded, and analyzed for their heat generation from the microcalorimeter is shown. First, *E. coli* BW25113 single-gene knockout cells (green), *E. coli* BW25113 wild-type (gray), and a negative control of media are seeded for overnight culture. Then, cells are inoculated at an initial OD_600_ value of 0.005 in a 20 mL glass ampoule. Ampoules are sealed and lowered into the microcalorimeter using ddH_2_O as the negative control. After 5h, the cells are removed from the microcalorimeter, and the final OD_600_ value is measured. The sample heat flow curves are analyzed for the peak value (*q*_*peak*_). This value is normalized by the final OD_600_ value to get a growth-normalized heat flow measurement.

### Protein-level regulators of thermogenesis

We probed the single-gene knockout space for increased normalized heat generation, which marked thermogenic dysregulation (Figure 2A and Table S1). Most strains fell within 10% of the wild-type *E. coli* BW25113 heat generation (Figure 2B). Only extremely high performers were considered in this experiment for further exploration. These strains exhibited an approximate 25% or greater increase in thermogenesis over wild type and were selected to be tested in biological triplicate. Most strains failed to maintain this increase threshold after triplicate measurement. However, through this analysis, three strains became evident as extreme heat producers. These are Δ*arcB* (*p* = 0.0031), Δ*glnL* (*p* = 0.0321), and Δ*yccC* (*p* = 0.0104). These strains had an average thermogenesis fold increase of 1.32 (Δ*glnL)*, 1.30 (Δ*yccC)*, and 1.25 (Δ*arcB)* over wild type. It is worth noting that in every instance, the thermogenesis of these strains was above the wild-type strain for the same experiment (Figure S1). Additionally, when tested using a student’s T-test and Welch’s correction, the increase was statistically significant (Figure 2C).

**Figure 2:**
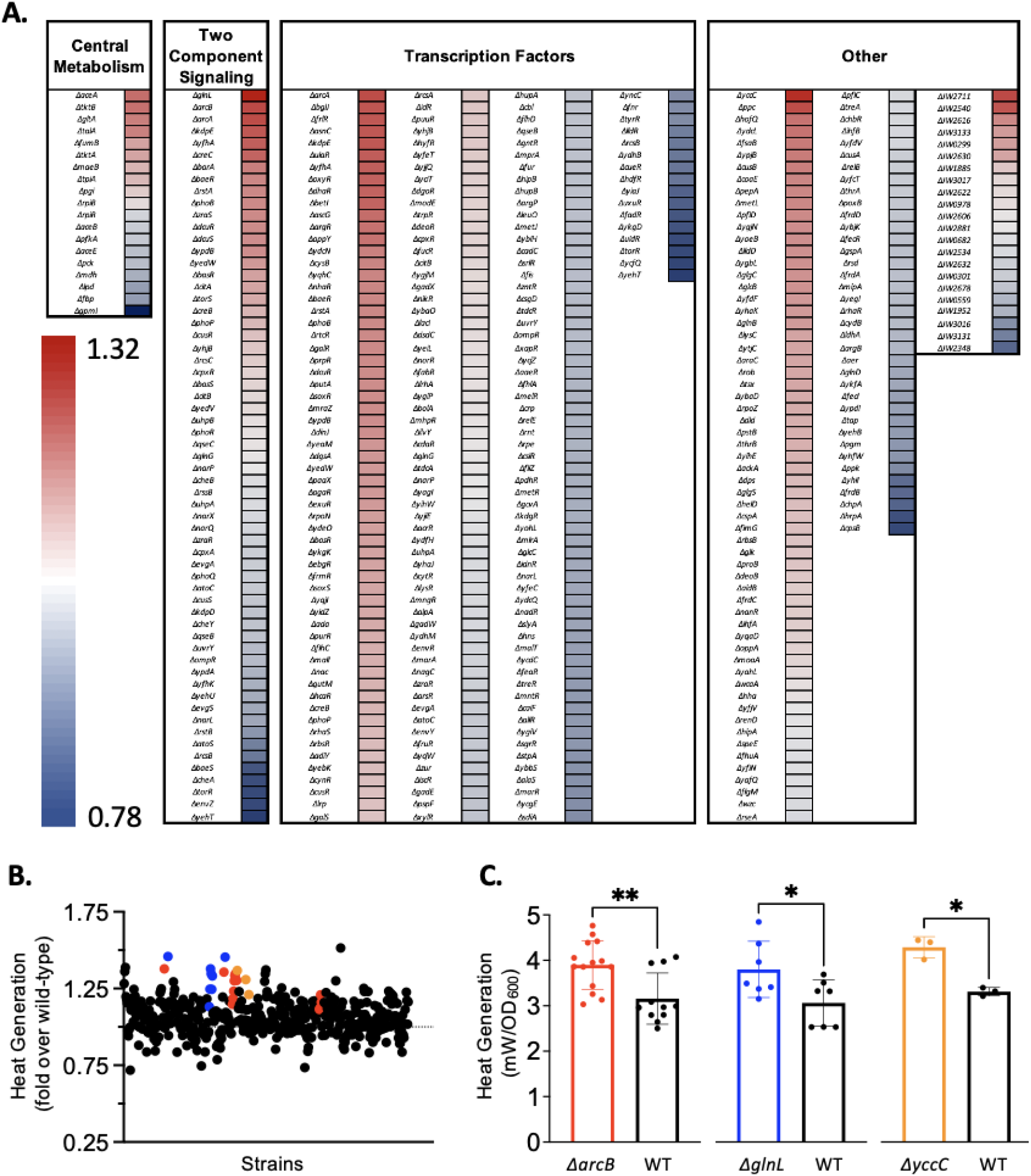
Heat generation screening of single-gene knockout strains. A) Heat map of categorized single-gene knockouts (heat flow fold change over wild-type; N = 449). Strains are categorized by their respective knockout gene’s function as central metabolism, two-component signal transduction, transcription factor, or other. “Other” strains either fall into none of the other categories or do not have 3 or 4 letter gene names assigned and are thus labeled by their JW ID. See Supplementary Table S1 for raw data. B) Scatter plot summarizing all results as fold change over wild-type. All tested single-gene knockout strains are shown (black). All data is normalized to wild-type result (dotted line) respective to experiment. C) Growth normalized heat flow (mW/OD_600_) bar graph against respective wild-type data (black). High-heat producers highlighted: *ΔarcB* (N = 14, t=3.335, and df=21.04) – red; *ΔglnL*(N = 8, t=2.435, and df=11.59) – blue; *ΔyccC* (N = 3, t=6.723, and df=2.599) – orange; and wild-type – black. Statistical significance by two-tailed T-test. *p<0.05; **p<0.01. Error bars: mean ± s.d.

Interestingly, these three genes encode proteins that are all autophosphorylating protein kinases. Specifically, *arcB* and *glnL* are well-studied global regulators of aerobic metabolism and nitrogen assimilation, respectively, and act through a two-component signal transduction pathway^48,49^. These are also two of the HKs with the highest phosphorylation rates^48^. Furthermore, since there are 29 HKs and 32 RRs in *E. coli*, some two-component signal transduction pathways have significant cross-talk between HK-RR pairs^47^. However, since we screened all the HK and RR genes individually, we minimized the impact of individual pairs’ crosstalk variations in our data.

Although the gene *yccC* (*etk*) is an autophosphorylating protein kinase, it is a tyrosine kinase that is not thought to be involved in two-component signaling in *E. coli*. Prokaryotic organisms are not thought to have sensor-protein tyrosine kinases, which are considered unique to eukaryotic organisms^50^. However, *etk* has been implicated in antibiotic resistance and is known to phosphorylate the heat shock sigma factor^51,52^.

Additionally, none of the transcription factor knockouts significantly increase cellular thermogenesis (Figure 2A). This result is unexpected as protein kinases generally regulate gene expression using transcription factors. Significantly increased thermogenesis in the protein kinase knockout mutants, but not the knockout mutants of the downstream transcription factor, can be explained in two ways. The knockout of a single protein kinase could increase the incidence of cross talk of HK-RR pairs. Alternatively, it is not the regulatory behavior of these protein kinases that is leading to these strains’ thermogenic effects, but instead the substrate-level effects of ATP that are causing this phenomenon of increased heat output. If the latter is the case, then the system would not only collapse to be more engineerable but also have a global regulator for cellular thermogenesis at the metabolite level, since all of the extreme thermogenic strains are the knockout mutants of self-phosphorylating protein kinases.

### Fundamental understanding of cellular thermogenesis at the chemical level

To better understand why a few strains increase in cellular thermogenesis while most do not, we wanted to explore some of the metabolic realities of extreme thermogenic cells. To do this, we moved forward with our three thermogenic strains: Δ*arcB*, Δ*glnL*, and Δ*yccC*. We hypothesized that the intracellular ATP concentration was the key metabolic regulator of this pathway. ATP is one of the highest energy compounds in the cell. It is generated from the catabolism of substrates to drive anabolic reactions^53^. Additionally, Ca^2+^-ATP coupled transporters have been implicated in mitochondrial thermogenesis in mammalian cells^26,54^.

We investigated intracellular ATP concentrations in our extreme thermogenic strains as well as a wild-type control over time. We correlated the ATP concentration alongside cellular growth (monitored by OD_600_) and heat flow output. The results show that intracellular ATP concentrations and cellular thermogenesis peak were at the same relative time of the growth curve, the late log phase (Figure 3). The intracellular ATP concentration could only be calculated after the 2h timepoint due to the sensitivity of the assay. The growth rates and curves of all strains are very similar (Figure 3A). Additionally, the general trends observed in the intracellular ATP concentration and the heat generation curves are conserved as well (Figure 3B, 3C).

**Figure 3:**
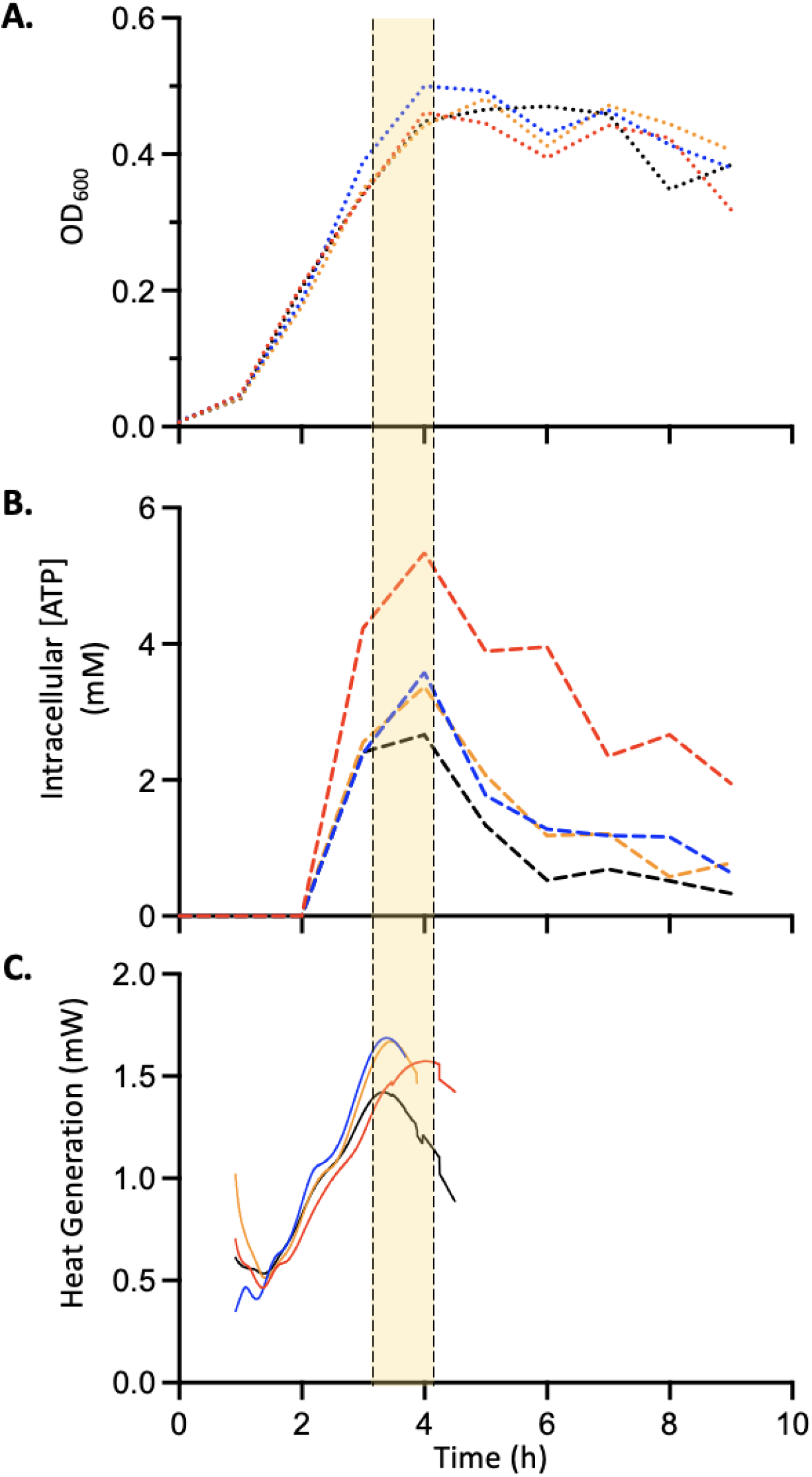
Microbial growth, intracellular ATP, and thermogenesis assay. A) Average strain growth (OD_600_) curves over time. B) Average strain intracellular ATP concentration (mM) over time C) Average strain heat flow (mW) curves over time. All samples were tested in triplicate, with distinct samples (N=3). *ΔarcB* – red, *ΔglnL* – blue, *ΔyccC* – orange, and wild-type – black.

During the late log phase, the cell is exiting the steady-state growth phase and is transitioning into the stationary phase. This transition is very dynamic and involves the coordinated downregulation and upregulation of proteins at the transcription, translation, and protein degradation level^55^. This massive shift in metabolism naturally leads to the accumulation of intracellular ATP. This result makes sense; when the ATP concentration is the highest, the cell has the most energy to spend, part of which is released as heat.

### Development of a microbial thermogenesis model

Hypothesizing that intracellular ATP is the metabolite that determines a cell’s thermogenesis, we should be able to extract the experimental thermogenesis data from experimentally derived parameters. Assuming that ATP is the sole, key metabolite responsible for cellular thermogenesis and that the dephosphorylation of ATP releases 7.3 kcal/mol, we can calculate the heat generation from a single cell and extend this to the entire population to model the cellular thermogenesis^56,57^. The inputs in this model are cellular growth and the intracellular ATP concentration, both as analytic functions of time. Cellular growth data was fit using the Gompertz growth equation, while the intracellular ATP concentration was fit using a lognormal distribution (Table S2). Then, using the constraint-based optimization (COBRA) model iML1515, the steady state ATP flux in an *E. coli* cell can be calculated^58^. This algorithm defines the Microbial Thermogenesis Model (MTM) (Figure 4A).

**Figure 4:**
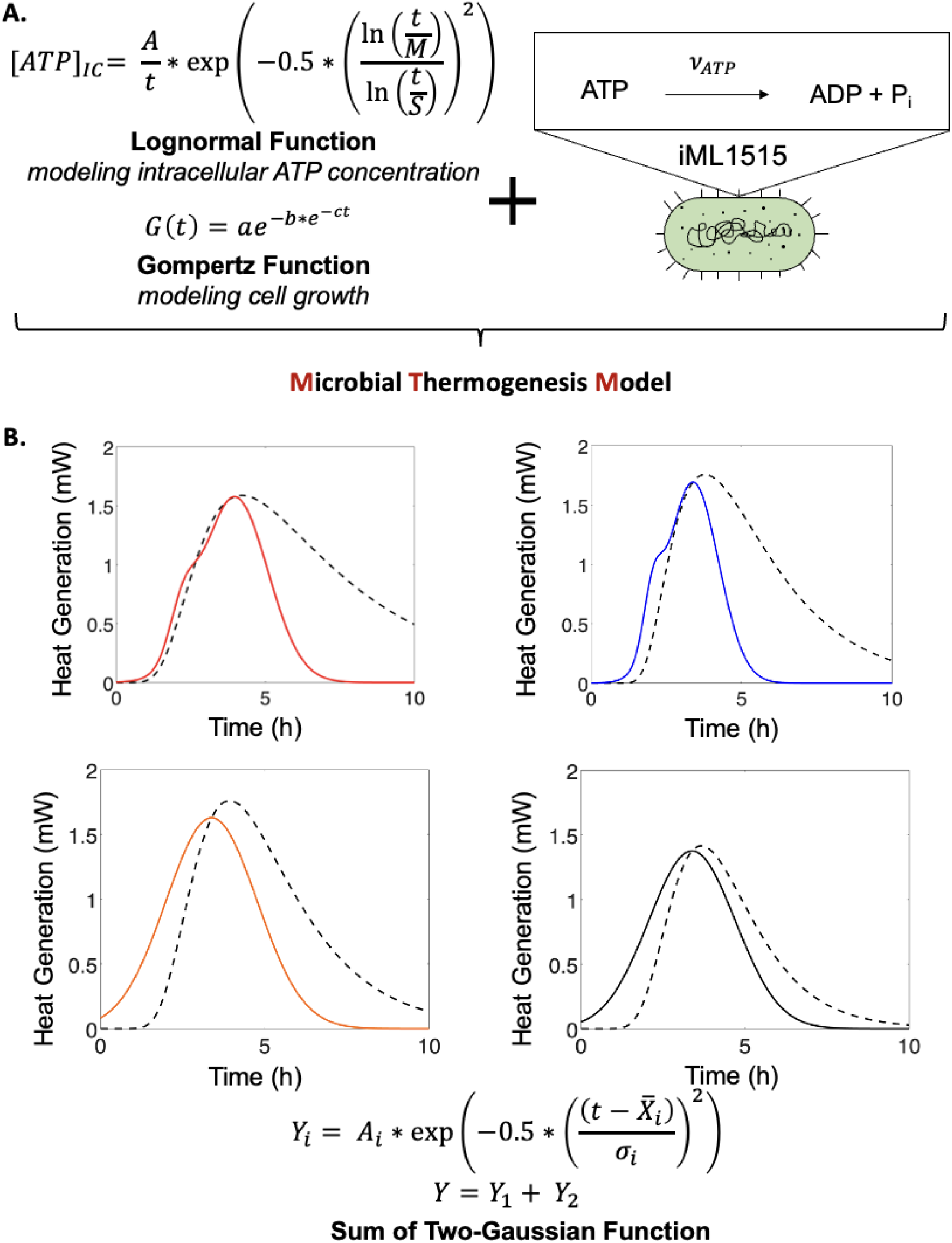
Microbial thermogenesis model (MTM) validation. A) MTM utilizes the Lognormal and Gompertz functions to model microbial intracellular ATP concentration and growth, respectively. Additionally, the model assumes an intracellular ATP pathway flux from wild-type *E. coli* calculated using iML1515. Gompertz growth equation: a – asymptote as *t* → ∞, b – displacement along x-axis, c – growth rate, and t – time. Lognormal equation: M – geometric mean, S – geometric standard deviation, A – factor related to amplitude. B) Average heat generation is fitted by the sum of two Gaussian equation (solid line) and plotted against the respective MTM approximations for each strain’s heat generation over time (dashed line). *ΔarcB* – red, *ΔglnL* – blue, *ΔyccC* – orange, and wild-type – black. Sum of two (*i =* 1, 2) Gaussian equation: A – amplitude, 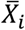 – mean, t – time, and *σ*_*i*_ – standard deviation.

The MTM matches well with the sum of two-Gaussian fit results. This is especially the case until the peak of cellular thermogenesis, when the correlation coefficients for all MTM fits are greater than 0.95. After this peak, the model deviates from the experimental results with correlation coefficients between 0.67 and 0.85 (Figure 4B and Table S3). This deviation is likely caused by the model’s assumptions. One assumption that leads to this deviation, at times greater than *t*_*peak*_, is that of steady state growth throughout the entire time domain of the model. This assumption is intrinsically accounted for in the MTM by the use of a single ATP flux calculated from the iML1515 model. The ATP flux must be maximal during steady state growth and decrease as the cell transitions its metabolism to the stationary phase. Nevertheless, the agreement between the experimental and model values supports the argument that there is a chemical-level control of cellular thermogenesis.

### Conclusion and future directions

Here, we present the protein kinases ArcB, GlnL, and YccC as the set of thermogenic regulators that are acting through intracellular ATP to modulate microbial thermogenesis. This fundamental study builds on previous work by extending it to engineer microbial systems for a variety of potential applications. One application is to develop probiotic strains with increased thermogenesis. The resulting increase in heat output by the gut microbiome would be the first step to developing a method for hypothermia prevention. Additionally, Irritable bowel syndrome (IBS), a disease known to affect 1 in 10 people globally, is often treated using heat therapy in the gut^59^. Building an environmentally safe, heat-generating, probiotic could lead to targeted heat therapy directly at the site of pathology for IBS treatments.

Although this work supports important conclusions in the field, there are some limitations. For example, the results cannot be directly extended to mammalian cells. Mammalian cells have distinct regulatory mechanisms that differ greatly from those found in microbial systems^60^. However, ATP is a metabolite whose importance transcends the boundaries of evolutionary domains. Thus, it is likely that this metabolite plays a key role in mammalian thermogenesis as well at the chemical level.

Further study is also needed to understand whether there are any metabolic enzymes, specifically those with ATP hydrolysis domains, which also regulate thermogenesis. Furthermore, there is immense value in understanding the exact biochemical pathways that influence cellular thermogenesis. This understanding is complicated by the delicate interactions of all proteins in signaling cascades such as the two-component pathways. Additionally, the complications compound as protein kinases are involved in a diverse set of sensing pathways and are intimately related to a cell’s environment. Understanding the role of these environmental signals in cellular heat generation will also become important as probiotics are developed. Furthermore, the human gut is anoxic^61^. Although some of these effects are captured in this study by using the gut-mimicking microaerobic environment to which cells are subjected, anoxemia still remains to be explored. Finally, the rational design of other highly thermogenic microbes will also proceed from this work.

## Methods

### Screening rationale and strain selection

A screen of single-gene knockouts of *Escherichia coli* BW25113 was conducted. A library of knockout strains, the Keio Collection, was purchased from Horizon Discovery (Waterbeach, United Kingdom)^46^. This library consists of single-gene knockouts of every non-essential gene in *E. coli* BW25113. This accounts for 3,985 genes out of the 4,453 genes present in the organism’s genome. The experiments were run in an 8-channel TAM Air Calorimeter (TA Instruments; New Castle, DE) using water as a negative control sample for each well, as is suggested for liquid samples in the instrument manual. For strain selection, a python script was developed to search the Keio Collection for genes of interest. See Supplementary Note 1 for details.

### Microbial thermogenesis assay

Six single-gene knockout strains were grown overnight along with one wild type (positive control) and one media control (negative control) for a total of 8 samples with shaking at 37°C and 250rpm in a 96-deep well plate. Samples are tested in singlicate (N = 1) for the initial screen. Then, if the sample presents >25% increase in thermogenesis, it was tested again (N ≥ 3). For all experiments, cells were grown in Luria–Bertani media (Sigma Aldrich 71753-6) supplemented with 2% glucose (Sigma Aldrich G7021). Following overnight incubation, OD_600_ (by Abs_600_, absorbance at 600 nm) was measured in a clear bottom 96-well plate, and the culture was diluted to a final OD_600_ of 0.005 in 10 mL growth media in a 20 mL glass ampoule (TA Instruments). The ampoule was then sealed and placed inside the TAM Air Calorimeter for 4-5h for heat flow measurement. After all samples had reached their peak, they were removed from the calorimeter and again measured for their OD_600_. See Supplementary Note 2 for more details.

### Thermogenesis data analysis

Initially, the data for each experiment were plotted as heat flow (W) over time (s). These data were baseline-normalized by subtracting out the negative control value from each sample value at each time point (collection frequency of 0.1Hz). Then, the data were normalized by final OD_600_ respective to the experimental strain to yield the normalized heat flow (mW/OD). To understand how the heat generation compares across experiments, the fold change was calculated by dividing each normalized heat flow by the wild-type normalized heat flow of that experiment. Final OD measurements were consistently taken within 30 min of the heat flow maximum. The calculation for significance was conducted by unpaired T-test, using Welch’s correction, using GraphPad Prism 9. All plots were also constructed using GraphPad Prism 9.

### Microaerobic bacterial growth assay

Cells were inoculated into growth media and cultured overnight from frozen stock stored at -80°C. Overnight culture was then seeded to a starting OD_600_ of 0.005 in 20 mL glass ampoules. Ampoules were capped and sealed at timepoint zero. Every 30 min, ampoules were uncapped, and Abs_600_ measurements were taken in a clear-bottomed, black-walled, 96-well plate. These measurement values were converted to OD_600_ by multiplying by 1.75, which was an experimentally determined conversion factor. All samples were tested in biological triplicate.

### Statistics

Statistical analyses and regressions were performed using Prism 9 (GraphPad). Quantitative values were averaged and shown by their means and standard deviations (s.d.). These values were tested for statistical significance of differences by student’s T-test using Welch’s correction to account for differences in sample’s standard deviations. These tests are two-tailed. Statistical significance is stated for samples with *p* values lower than 0.05. For the model quantification, correlation coefficients were calculated to test for model fit to data. Correlation coefficients were calculated using MATLAB 2022a. Coefficients closer to 1.0 reveal a higher coincidence between two samples.

### ATP quantification assay

ATP measurements were conducted in the following way. Cells were grown in 20 mL glass ampoules (TA instruments) in 10 mL of growth media. The ampoules were capped and sealed at time point zero. Every 30 min, ampoules were unsealed, and the intracellular ATP concentration was measured in triplicate for each strain (Δ*arcB*, Δ*glnL*, Δ*yccC*, and wild-type) as follows. The cells were first lysed, using boiling water prepared on a magnetic hot plate. From the ampoule, 1 mL of the sample was taken into a 1.5 mL centrifuge tube (containing about 10^6^ cells) and pelleted in a microcentrifuge at 12,000xg for 10 min. A luciferin-luciferase luminescent reporter assay was then used to quantify the intracellular ATP concentration (Invitrogen). OD_600_ measurements were also made at each time-point and used to calculate an average intracellular ATP concentration with 1 OD_600_ as 10^9^ cells/mL and an average cell volume as 1fL^62,63^.

### Microbial thermogenesis model construction

The microbial thermogenesis model (MTM) was developed under three key assumptions: the sole metabolic determinant of microbial thermogenesis is ATP; the intracellular ATP flux scales with the intracellular ATP concentration; the ATP flux is constant at its steady-state value throughout the thermogenesis assay. The growth, the intracellular ATP concentration, and thermogenesis raw data were fit using the Gompertz growth, lognormal, and sum of two Gaussian equations, respectively, to get analytic functions over time representing the data (Figure S2-4). All models demonstrated good agreement with raw data (Table S2). Model fitting and correlation coefficient calculations were conducted using Microsoft Excel and GraphPad Prism 9. A MATLAB algorithm was then developed to output a heat generation prediction (mW) from input growth, ATP concentration, and ATP flux parameters. The steady-state ATP metabolic flux parameters were calculated in a python script using constraint-based optimization of metabolite flux from the COBRApy package and iML1515 model. In short, the MATLAB algorithm converts growth data into cell number data, which were then used to calculate the total intracellular ATP concentration-weighted ATP flux through each cell. Then, this flux was converted to a heat energy release for the entire culture.

## Data Availability

Raw calorimetric and growth data are available upon request. Gene annotations are available through Metacyc (https://metacyc.org/).

## Code Availability

Model development was conducted in MATLAB. All codes for the generation of the MTM model are available on the Microbial Thermogenesis Model GitHub repository: (https://github.com/pdhatt/microbialthermogenesismodel.git).

## Acknowledgements

The authors thank Dr. Jinjin Diao for his comments and feedback on the manuscript. This work was supported by the Office of Naval Research (N00014-21-1-2206 to T.S.M). The content is solely the responsibility of the authors and does not necessarily represent the official views of the funding agency.

## Author contributions

T.S.M. conceived the project. P.S.D. performed the experiments. P.S.D. and T.S.M. designed experiments, analyzed the data, and wrote the manuscript.

